## Supplemental Table 3 for "Olfactory ensheathing cells from adult female rats are hybrid glia that promote neural repair" (from another version of this preprint: version 2, 2024-09-25)

### Supplemental Table 3: Top 20 marker genes based on log2 fold change in each of the 5 OEC clusters revealed by scRNA-seq

#### **Cluster 0 –** Genes listed in descending order

| OEC sub-cluster genes (Fold change) | Name and Function | Where is it expressed? | References |
|---|---|---|---|
| **1** *Pmp22* (1.427) | Peripheral myelin protein 22 is a myelin associated gene | Schwann cell myelin | (Notterpek et al., 1999; Roet et al., 2011; Salzer, 2015) |
| **2** *Ccdc80* (1.166) | Coiled-coil Domain containing 80, secreted, part of Hedgehog pathway, may regulate vascular function | Zebrafish somites, axonal pathfinding | (Brusegan et al., 2012; Della Noce et al., 2015; Sasagawa et al., 2016) |
| **3** *Gldn* (1.130) | Gliomedin is a required ECM protein that initiates assembly of PNS nodes of Ranvier | Schwann cell surface | (Colognato & Tzvetanova, 2011; Eshed et al., 2005) |
| **4** *Ccn3/Nov* (1.165) | Nephroblastoma over-expressed, matricellular protein with proregenerative function, Ccn3 upregulates chemokines CCL2 and CXCL1 | Secreted by astrocytes & regulatory T cells; promotes oligodendro-cyte differentiation and myelination | (de la Vega Gallardo et al., 2020; Le Dreau et al., 2010; Malik et al., 2015) |
| **5** *Col11A1* (1.049) | Collagen, type XI, alpha 1 ECM protein | Schwann cells | (Zhang et al., 2021) |
| **6** *Serpinf1* 1.035) | Serine protease inhibitor, Neurotrophic functions, enhances axon outgrowth | Dorsal root ganglion axons *in vitro* | (Roet et al., 2011; Roet et al., 2013) |
| **7** *Fbln2* (1.026) | Fibulin2; mediates the TGF-β1 signaling in adult neural stem cells | Astrocytes in adult neurogenesis (SVZ) | (Radice et al., 2015) |
| **8** *Nefm* (0.976) | Neurofilament medium | Glia, neurons | (Roet et al., 2011) |
| **9** *Ccn2* (0.91) | Connective tissue growth factor, secreted CTGF regulates growth factors, cytokines and MMPs | Astrocytes, Schwann cells | (Jun & Lau, 2011; Lamond & Barnett, 2013; Malik et al., 2015; Roet et al., 2011) |
| **10***RGD1305645* (0.87) | *Ecrg4* augurin precursor, esophageal cancer-related gene 4 protein | Uncharacterized in nervous system | Rat Genome Database https://rgd.mcw.edu |
| **11** *Nqo1* (0.849) | NAD(P)H dehydrogenase, quinone 1; protect CNS cells against oxidative stress | Bergmann glia, astrocytes, oligodendrocytes | (Roet et al., 2011; Stringer et al., 2004) |
| **12** *Marcks* (0.839) | Myristoylated alanine-rich C-kinase substrate protein; actin-cross-linking protein regulates radial glia function | Astrocytes, oligodendrocytes | (Artegiani et al., 2017; Roet et al., 2011; Weimer et al., 2009) |
| **13** *Prss23* (0.814) | Novel vascular protease required for endothelial-to-mesenchymal transition | Cardiovascular valve development | (Chen et al., 2013; Roet et al., 2011) |
| **14** *Rarres2* (0.791) | Chemerin or Retinoic acid receptor responder 2, an inflammatory chemokine; under neuroendocrine control | Ventricular ependymal cells, tanycytes | (Helfer et al., 2016) |
| **15** *Cck* (0.775) | Cholecystokinin, gut hormone and neuropeptide | Peripheral nerves, neurons | (Rehfeld, 2017) |
| **16** *Fst* (0.767) | Follistatin; plays a role in myelination and controls activin | Olfactory bulb, ONL; Areas of adult neurogenesis, | (Ogawa et al., 2020) |

| | | | |
|---|---|---|---|
| **17** *Tagln* (0.767) | Transgelin, a TGFβ-inducible gene that functions as an actin-crosslinking protein | Vascular endothelial, smooth muscle, neural crest stem cells | (Elsafadi et al., 2016; Roet et al., 2011) |
| **18** *Fn1* (0.728) | Fibronectin-1; major component of ECM | Perivascular fibro-blasts, macrophages, reactive astrocytes | (Colognato & Tzvetanova, 2011; Zhu et al., 2015) |
| **19** *Cd200* (0.728*)* | CD200 binds to and activates its receptor (CD200R) on microglial cells, have immunoregulatory functions | Neurons, astrocytes, oligodendrocytes | (Manich et al., 2019) |
| **20** *Dpysl3* (0.691) | Dihydropyrimidinase-like 3; important for microglia activation, migration and phagocytic ability | Ameboid microglia | (Manivannan et al., 2013) |

### **Cluster 1–** Genes listed in descending order

| **OEC sub-cluster genes (Fold change)** | **Name and Function** | **Where is it expressed?** | **References** |
|---|---|---|---|
| **1** *Pcsk1* (1.616) | Preprohormone convertase 3 or Neuroendocrine convertase 1 | Schwann cells | (Marcinkiewicz et al., 1998; Roet et al., 2011) |
| **2** *Gal* (1.608) | Galanin, small neuropeptide member of corticotropin-releasing factor family; regulates oligodendrocyte survival | Neurons, neural progenitor cells in areas of adult neurogenesis, oligodendrocytes | (Cordero-Llana et al., 2014; Gresle et al., 2015; Roet et al., 2011; Ubink et al., 2003) |
| **3** *Atf3* (1.234) | Activating transcription factor 3, a pro-regenerative intrinsic factor enhances PNS axon regeneration | Schwann cells, injured dorsal root ganglia, axotomized axons | (Hunt et al., 2004; Mahar & Cavalli, 2018; Shortland et al., 2006; Zhang et al., 2021) |
| **4** *Ucn2* (1.223) | Urocortin-2 protein member, neuropeptide involved in stress response and axonal outgrowth, corticotropin-releasing factor family | Hypothalamic neurons, motor neurons in brain stem, perisynaptic Schwann cells | (D'Este et al., 2022; Reyes et al., 2001; Roet et al., 2011) |
| **5** *Crabp2* (1.142) | Cellular retinoic binding protein 2 transports retinoic acid to the nuclear receptors | Neural crest-derived cranial ganglia, olfactory placode, olfactory epithelium | (Cai et al., 2012) |
| **6** *Gap43* (1.137) | Growth associated protein 43, transiently expressed in growing axons and after axotomy | Growing axons, Schwann cell precursors, | (Jacobson et al., 1986; Sensenbrenner et al., 1997; Takeoka et al., 2011) |
| **7** *Btc* (1.079) | Betacellulin, part of Epidermal Growth Factor family, a ligand for EGFR and a growth factor | Myelinating Schwann cells | (Roet et al., 2011; Vallieres et al., 2017) |
| **8** *LOC108348064* (1.078) | Aph-1 homolog B, gamma secretase subunit, participates in Notch pathway | Predicted location at synapse and transport vesicle | Rat Genome Database https://rgd.mcw.edu |
| **9** *Chrdl2* (1.066) | Chordin-like 2, a novel secreted BMP binding inhibitor | Chondrocytes, regulates dorso-ventral patterning | (Branam et al., 2010; Nakayama et al., 2004) |
| **10** *Cast* (1.038) | Calpastatin, endogenous calpain inhibitor, key regulator of axon survival post injury and during development | Broadly expressed in CNS & PNS, including Purkinje neurons and retinal ganglion cells. | (Yang et al., 2013) |
| **11** *Il11* (0.961) | Interleukin-11, secreted by astrocytes, enhances oligodendrocyte survival, maturation and myelin formation | Astrocytes | (Roet et al., 2011; Zhang et al., 2006) |

| | | | |
|---|---|---|---|
| **12** *Ngfr* (0.96) | Nerve growth receptor-p75 enhances axonal growth and remyelination | Schwann cells | (Roet et al., 2011; Tomita et al., 2007) |
| **13** *Nes* (0.953) | Nestin, an intermediate filament associated with neural stem cells | Neuroepithelial stem cells, radial glia, adult neural crest-related stem cells | (Cattaneo & McKay, 1990; Widera et al., 2011) |
| **14** *S100β* (0.918) | S100β is a calcium binding protein that increases after injury and supports neurogenesis | Schwann cells, astrocytes, NG2 cells, APC+ oligodendrocytes in olfactory bulb | (Su et al., 2021; Villarreal et al., 2014) |
| **15** *Vcan* (0.903) | Versican, a chondroitin sulfate proteoglycan | Astrocytes, late oligodendrocyte progenitors | (Anderson et al., 2016; Perlman et al., 2020) |
| **16** *Bdnf* (0.901) | Brain-derived neurotrophic factor, axonal growth and regeneration, dendritic growth | Schwann cells | (Gordon, 2009; Guerout et al., 2010; Meyer et al., 1992; Runyan & Phelps, 2009) |
| **17** *Stim1* (0.884) | Stromal interaction molecule, $Ca^{2+}$ sensor in ER that mediates store-operated $Ca^{2+}$ entry (SOCE) | Astrocytes, microglia | (Michaelis et al., 2015; Muller et al., 2014) |
| **18** *Serpinb6a* (0.874) | Serpin family B member 6a, part of serine protease inhibitor complex, loss causes hearing deficit | Auditory related neurosensory epithelium | Rat Genome Database https://rgd.mcw.edu |
| **19** *Hspb1* (0.858) | Heat shock protein β1, small heat shock protein | Highly expressed in glial cells | (Lopez-Gonzalez et al., 2014) |
| **20** *LOC102549-726* (0.811) | Uncharacterized | Uncharacterized | Rat Genome Database https://rgd.mcw.edu |

### **Cluster 2 –** Genes listed in descending order

| **OEC sub-cluster genes (Fold change)** | **Name and Function** | **Where is it expressed in progenitor cells?** | **References** |
|---|---|---|---|
| **1** *Stmn1* (2.502) | Stathmin 1, microtubule destabilizing protein antagonizes differentiation of oligodendrocytes | Areas of adult neurogenesis-SVZ, Oligodendrocyte progenitors, astrocytes | (Boekhoorn et al., 2014; Liu et al., 2003) |
| **2** *Cenpf* (2.265) | Centromere protein F, associates with centromere-kinetochore complex | In nuclear matrix during G2 phase in interphase, Schwann cells | (Whitfield et al., 2002; Zhang et al., 2021) |
| **3** *Prc1* (2.118) | Polycomb repressive complex 1, repressive epigenetic mechanism that controls stem cell self-renewal or differentiation | Widely expressed in neurons and glia, proliferative globose basal cells in olfactory epithelium | (Goldstein et al., 2018) |
| **4** *Hmgb2* (2.099) | High mobility group box 2, secreted proinflammatory cytokine | Expressed in the nervous system during embryogenesis | (Lee et al., 2014) |
| **5** *Pttg1* (1.971) | Pituitary tumor transforming gene 1 | Regulates cell cycle M/G1, sister chromatid cohesion | (Whitfield et al., 2002) |
| **6** *Ube2c* (1.966) | Ubiquitin conjugating enzyme E2 C | Regulates cell cycle G2/M phase | (Clark et al., 2019; Whitfield et al., 2002) |
| **7** *Cdk1* (1.951) | Cyclin dependent kinase 1 binds to Cyclin B1 to enter mitosis | Regulation of cell cycle, mitosis | (Martinsson-Ahlzen et al., 2008; Morgan, 1997) |

| | | | |
|---|---|---|---|
| **8** *Cenpw* (1.895) | Component of CENPA-Nucleosome-associated complex required in progenitor proliferation | Required during mitosis for kinetochore formation and centriole splitting | (Aygun et al., 2021) |
| **9** *Cdkn3* (1.876) | Cyclin dependent kinase inhibitor 3 interacts with Cdk2 and prevents its activation | Cell cycle regulation, G1/S transition | (Nalepa et al., 2013) |
| **10** *Top2a* (1.855) | DNA topoisomerase II alpha | Cell cycle regulation, peaks in G2 | (Whitfield et al., 2002) |
| **11** *Tpx2* (1.855) | TPX2 microtubule nucleation factor, part of acentrosomal MT organizing center | Localized to centrosome and neurite shaft | (Chen et al., 2017) |
| **12** *Cks2* (1.828) | CDC28 protein kinase regulatory subunit 2, counteracts Cks1 and stabilizes p27 | Active from late G1 to mitosis, regulator of stem cell maintenance and differentiation | (Frontini et al., 2012; Martinsson-Ahlzen et al., 2008) |
| **13***LOC100359539* (1.808) | Ribonucleotide reductase M2 polypeptide | Involved in deoxyribonucleotide biosynthetic process | Rat Genome Database https://rgd.mcw.edu |
| **14** *Spc24* (1.706) | SPC24 component of NDC80 kinetochore complex | Kinetochore-microtubule attachments and spindle checkpoint activity | (McCleland et al., 2004) |
| **15** *Dut* (1.614) | Deoxyuridine triphosphatase (dUTPase), prevents uracil misincorporation into DNA | Essential enzyme for nucleotide metabolism and DNA repair | (Ye et al., 2019) |
| **16** *Mki67* (1.613) | Ki-67 is an early progenitor marker of proliferation, component of mitotic cell periphery | Nuclei of proliferating cells, Schwann cell sub-type 4 | (Artegiani et al., 2017; Zhang et al., 2021) |
| **17***LOC100360316 Hmgn2-ps4* (1.585) | High mobility group nucleosomal binding domain 2 | Chromatin in the nucleus | Rat Genome Database https://rgd.mcw.edu |
| **18** *Ckap2* (1.573) | Cytoskeleton associated protein 2, stabilizes microtubules | Maintains microtubule nucleation sites, regulates cell division | (Case et al., 2013) |
| **19** *Fam111a* (1.512) | Family with sequence similarity 111member A, cell-cycle regulated | Mediates effects of protein obstacles on DNA replication forks | (Kojima et al., 2020) |
| **20** *Cdca8* (1.493) | Cell division cycle associated 8, part of a chromosomal passenger complex | Required for chromatin-induced microtubule stabilization and spindle formation | (Sampath et al., 2004) |

### **Cluster 3 –** Genes listed in descending order

| **OEC sub-cluster genes (Fold change)** | **Name and Function** | **Where is it expressed?** | **References** |
|---|---|---|---|
| **1** *C1qa* (3.981) | Complement C1q subcomponent a, part of the innate immune system | Macrophages, microglia | (Durante et al., 2020; Zhang et al., 2014) |
| **2** *Tyrobp* (3.944) | TYRO protein tyrosine kinase binding protein (DAP 12), a microglial membrane adaptor that mediates microglial upregulation and neuropathic pain | Microglia | (Guan et al., 2016; Zhang et al., 2014) |
| **3** *C1qb* | Complement C1q subcomponent b | Macrophages, microglia | (Durante et al., 2020; Zhang et |

| | | | |
|---|---|---|---|
| (3.905) | | | al., 2014) |
| **4** *Apoe* **(**3.877) | Apolipoprotein E is a cholesterol transport gene with many functions, binds to low-density lipoprotein receptor gene family | Astrocytes, cultured microglia | (Cadiz et al., 2022; Herz & Chen, 2006) |
| **5** *Fcer1g* (3.665) | Fc receptor, IgE, high-affinity I, gamma polypeptide. Activates microglial phagocytosis pathways | Microglia, astrocytes, oligodendrocytes, neurons | (Sierksma et al., 2020; Zhang et al., 2014) |
| **6** *Cst3* (3.499) | Cystatin C protein, | Microglia | (Zhang et al., 2014) |
| **7** *Tmem176b* (3.491) | Transmembrane protein 176b | Immature monocytes and dendritic cells, Schwann cells | (Condamine et al., 2010; Zhang et al., 2021) |
| **8** *Anxa3* (3.4) | Annexin A3, calcium-dependent lipid binding protein, | Microglia, monocytes | (Smithson & Kawaja, 2010) |
| **9** *Tmem176a* (3.365) | Transmembrane protein 176, paralog of Tmem176b | Immature monocytes and dendritic cells | (Condamine et al., 2010) |
| **10** *C1qc* (3.183) | Complement C1q subcomponent c | Macrophages, microglia | (Cadiz et al., 2022; Durante et al., 2020; Zhang et al., 2014) |
| **11** *Aif1* (3.033) | Allograft inflammatory factor 1, ionized calcium-binding adaptor molecule (Iba1), marker to identify microglia | Macrophages, microglia | (Artegiani et al., 2017) |
| **12** *Csf1r* (2.559) | Colony stimulating factor 1 receptor, binds to colony stimulating factor 1, is upregulated after injury | Surface of microglia | (Guan et al., 2016; Roet et al., 2011; Zhang et al., 2014) |
| **13** *Ctsz* (2.436) | Cathepsin Z, lysosomal cysteine proteinase, enables cysteine-type endopeptidase activity | Microglia | (Zhang et al., 2014) Rat Genome Database |
| **14** *Laptm5* (2.338) | Lysosomal protein transmembrane 5 | Microglia | (Artegiani et al., 2017; Roet et al., 2011) |
| **15** *Ctsb* (2.288) | Cathepsin B, lysosomal cysteine proteinase involved in phagocytosis | Microglia | (Grubman et al., 2021; Zhang et al., 2014) |
| **16** *Lyz2* (2.262) | Lysozyme C type 2, lysosomal gene | Microglia | (Cadiz et al., 2022; Roet et al., 2011) |
| **17** *Rgs10* (2.237) | Regulator of G protein signaling 10, in RGS superfamily, selectively interacts with Gαi proteins | Microglia, monocyte-derived dendritic cells | (Almutairi et al., 2020; Artegiani et al., 2017) |
| **18** *Cd68* (2.134) | Cluster of Differentiation 68, glycoprotein associated with endosomal/lysosomal compartment | Microglia, Macrophages, other mononuclear phagocytes | (Chistiakov et al., 2017) |
| **19** *Arhgdib* (1.993) | Rho GDP dissociation inhibitor-β | Microglia | (Artegiani et al., 2017) |
| **20** *Trem2* (1.908) | Triggering receptor expressed on myeloid cells 2, Trem2 protein interacts with protein produced from *Tyrobp* gene | Microglia, immature dendritic cells | (Artegiani et al., 2017; Zhang et al., 2014) |

**Cluster 4 –** Genes listed in descending order

| **OEC sub-cluster genes (Fold change)** | **Name and Function** | **Where is it expressed?** | **References** |
|---|---|---|---|
| **1** *Sidt2* (2.833) | SID1 transmembrane family member 2 binds RNA/DNA, transports nucleic acids to lysosomes for degradation | Brain lysosomes | (Hase et al., 2020; Jialin et al., 2010) |
| **2** *Ckb* (2.687) | Brain-type creatine kinase, fuels ATP-dependent cytoskeletal dynamics, facilitates spreading of cortical actin for motility | Astrocytes, oligodendrocytes, macrophages | (Kuiper et al., 2009) |
| **3** *Npvf* (2.196) | Neuropeptide VF precursor or gonadotropin-inhibitory hormone. | Hypothalamic neurons, gonads, olfactory bulbs-terminal nerve | (Ubuka et al., 2016) |
| **4** *Ptn* (2.127) | Pleiotrophin, secreted-heparin-binding protein, neurotrophic growth factor for neurite and glial process outgrowth during development and after injury | Microglia, macrophages, astrocytes, pericytes, endothelial cells | (Nikolakopoulou et al., 2019; Yeh et al., 1998) |
| **5** *C1ql1* (1.928) | Complement C1q like 1, involved in different aspects of synaptic remodeling, synapse elimination on Purkinje cells | Neurons such as cerebellar granule cells, oligodendrocyte precursors | (Eroglu & Barres, 2010; Kakegawa et al., 2015; Zhang et al., 2014) |
| **6** *Igfbp3* (1.924) | Insulin-like growth-factor binding protein-3 reduces apoptosis of pericytes and retinal neurons | Astrocytes, endothelial cells | (Kielczewski et al., 2011) (Watanabe et al., 2015) |
| **7** *Gria2* (1.922) | Glutamate receptor ionotropic AMPA 2 determines channel conductance and $Ca^{2+}$ permeabi-lity, with RNA editing (Q/R site) channel is impermeable to $Ca^{2+}$ | Astrocytes, Oligodendrocyte precursors (OPCs) | (Chen et al., 2018; Rusnakova et al., 2013; Zhang et al., 2014) |
| **8** *Stmn2* (1.842) | Stathmin2 is a tubulin-binding protein that regulates microtubule disassembly | Neural stem cells, neural crest cells, neurons | (He et al., 2016; Himi et al., 1994; Zhang et al., 2014) |
| **9** *Actn1* (1.825) | Actinin, alpha1, cross-links actin filaments to maintain cell scaffolds & mechanical stability | Migrating cells | (Sjoblom et al., 2008) |
| **10** *Vgf* (1.783) | Secreted nerve growth factor with neuroendocrine functions, promotes oligodendrogenesis | Neurons and neuroendocrine cells | (Alvarez-Saavedra et al., 2016; Zhang et al., 2014) |
| **11** *Col8a1* (1.646) | Collagen, type 8, alpha1, short chain collagen involved in cell adhesion, migration, & sprouting angiogenesis | Endothelial cells, vasculature, smooth muscle, mast cells | (Illidge et al., 1998) |
| **12** *Nacc2* (1.537) | Nacc family member 2, multiple functions such as DNA-binding repressor | Chromatin, nucleolus, mitochondrion | Rat Genome Database https://rgd.mcw.edu |
| **13** *Thbs2* (1.472) | Thrombospondin 2, secreted ECM glycoprotein contributes to synaptogenesis during development and repair of blood-brain barrier | astrocytes | (Christopherson et al., 2005; Tian et al., 2011) |
| **14** *Bgn* (1.465) | Biglycan, secreted chondroitin sulfate proteoglycan with neuronal survival-enhancing | Meningeal fibroblasts, pericytes | (Junghans et al., 1995; Zhang et al., 2014) |

| | | | |
|---|---|---|---|
| | activity | | |
| **15** *Mt3* (1.462) | Metallothionein-3, maintains homeostasis of copper and zinc, regulates lysosomal function, associated with growth inhibitory factor | Brain, astrocytes | (Lee et al., 2010) |
| **16** *Rarres1* (1.458) | Retinoic acid receptor responder 1. Expression induced by retinoic acid, mainly studied in cancer as a potential tumor suppressor | Subset of stellate and superior cervical sympathetic ganglion neurons (NA1) | (Furlan et al., 2016; Mapps et al., 2022; Roet et al., 2011) |
| **17** *Tubb2b* (1.458) | Tubulin β, class 2b, together with α-tubulin form microtubules critical for cell movement or neuronal migration | Neurons, radial glia | (Chakraborti et al., 2016) |
| **18** *Tubb3* (1.435) | Tubulin β, class 3, neuronal specific, together with α-tubulin form microtubules critical for migration axonal guidance and transport | Neurons, dorsal root ganglion | (Chakraborti et al., 2016; Zhang et al., 2021; Zhang et al., 2014) |
| **19** *C1qtnf5* (1.413) | C1q tumor necrosis factor 5, secreted membrane-associated protein in RPE, when mutated, destabilizes protein & leads to photoreceptor death | Retinal pigment epithelium (RPE) | (Stanton et al., 2017) |
| **20** *Gpm6b* (1.345) | Glycoprotein M6b, proteolipid protein family member binds cholesterol and contributes to CNS myelin, in Schwann cell nodes of Ranvier | Oligodendroglia, Schwann cell microvilli, neurons, | (Bang et al., 2018; Roet et al., 2011; Werner et al., 2013) |
