## Supplemental Table 4 for "Olfactory ensheathing cells from adult female rats are hybrid glia that promote neural repair" (from another version of this preprint: version 2, 2024-09-25)

### Supplemental Table 4: Primary antibodies used in study with working dilutions, assays, and sources

| Primary Antisera, Host Species | Immunogen | Source; Catalogue # RRID | Working Dilutions, IHC |
|---|---|---|---|
| Activating transcription factor 3 (Atf3), Mouse monoclonal | Recombinant fragment corresponding to Human Atf3 aa 1-150 | Abcam; #ab191513 RRID:AB_2868437 | 1:4000; TSA + Citric Acid |
| Allograft inflammatory factor 1/ Ionized calcium-binding adaptor molecule 1 (Aif1/Iba1), Rabbit polyclonal | Recombinant protein within the center region of human Iba1 | Wako (Osaka, Japan); #019-19741 RRID:AB_839504 | 1:5000; IF |
| Annexin A3 (Anxa3), Rabbit polyclonal | Recombinant protein within the center region of human Annexin III | Gene Tex (USA); #GTX103330 RRID:AB_11174712 | 1:5000; TSA + Citric Acid |
| Apolipoprotein E (Apoe), Goat **p**olyclonal | Recombinant human apolipoprotein E | Sigma-Aldrich; #AB947 RRID:AB_2258475 | 1:2000; TSA + Citric Acid |
| Brain lipid-binding protein (Blbp), Rabbit monoclonal | GST/His-tagged full-length recombinant human Brain lipid-binding protein | Sigma-Aldrich; #ZRB13190 RRID:AB_2920766 | 1:5000; ICC, IF |
| Brain lipid-binding protein (Blbp), Rabbit polyclonal | GST-tagged recombinant protein corresponding to human Brain lipid-binding protein | Sigma-Aldrich; #ABN14 RRID:AB_10000325 | 1:1500; ICC, IF |
| Gliomedin (Gldn), Rabbit polyclonal | Synthetic peptide | Abcam (Cambridge, MA); #ab24483 RRID:AB_2111616 | 1:100; IF + Citric Acid |
| Growth-associated protein (Gap43), Rabbit polyclonal | Synthetic peptide corresponding to Rat GAP43 aa 200 to the C-terminus conjugated to keyhole limpet haemocyanin | Abcam (Cambridge, MA); #ab16053 RRID:AB_443303 | 1:500; IF |
| Green fluorescent protein (GFP), Chicken polyclonal | Recombinant GFP expressed in Escherichia coli | Aves Labs Inc.; #GFP-1020 RRID:AB_10000240 | 1:1000; ICC, IF |
| Glial fibrillary acidic protein (GFAP), Mouse monoclonal | Cow spinal cord homogenate | BD Biosciences; #556327 RRID:AB_396365 | 1000; IF |
| Glyceraldehyde-3-phosphate dehydrogenase (GAPDH), Mouse monoclonal | Glyceraldehyde-3-phosphate dehydrogenase from rabbit muscle | Sigma-Aldrich; #MAB374 RRID:AB_2107445 | 1:100K; WB |
| L1, Rabbit polyclonal | Affinity purified L1 from rat plasma membranes | Gift from Dr. Vance Lemmon | 1:10,000; ICC |
| Marker of proliferation ki-67 (Mki67), Rabbit polyclonal | Synthetic peptide | Abcam (Cambridge, MA); #ab15580 RRID:AB_443209 | 1:500; ICC<br>1:5000; TSA |
| Nestin (Nes), Mouse monoclonal | Spinal cord homogenate | Developmental Studies Hybridoma Bank; #rat-401 RRID:AB_2235915 | 1:1000; TSA + Citric Acid |
| Neurofilament-200 (NF), Rabbit polyclonal | Recombinant fusion protein containing the extreme C-terminus of rat NF-H | Sigma-Aldrich; #AB1989 RRID:AB_91202 | 1000; IF |
| N-Cadherins, Mouse monoclonal | Recombinant domain corresponding to the intracellular domain of chicken N-Cadherin | Invitrogen; #33-3900 RRID:AB_2313779 | 1:500; ICC |
| Neural cell adhesion molecule (NCAM), Rabbit polyclonal | Highly purified chicken NCAM | Sigma-Aldrich; #AB5032 RRID:AB_2291692 | 1:5000; ICC |
| Nerve growth factor receptor, p75 ($Ngfr^{p75}$), Mouse monoclonal | Hybridoma line 192 obtained by fusing Sp2/0- Ag14 myeloma cells with splenocytes from Balb/C mice | Protocol adapted from Chandler et al., 1984 | 1:50; ICC<br>1:5; Immunopurification |
| Nerve growth factor receptor, p75 ($Ngfr^{p75}$), Rabbit polyclonal | Recombinant mouse NGF Receptor p75 extracellular fragment | Sigma-Aldrich; #AB1554 RRID:AB_90760 | 1:30K; ICC<br>1:1500; Immunopurification |
| Peripheral myelin protein 22 (Pmp22), Rabbit polyclonal | Synthetic peptide from human Pmp22 | Gene Tex (USA); #GTX85834 RRID:AB_10733560 | 1:1000; TSA + MeOH* |
| Reelin (Reln), Goat polyclonal | Mouse myeloma cell line NS0-derived | R&D Systems; #AF3820 | 1:250; IF |

| | | | |
|---|---|---|---|
| | recombinant mouse Reelin | RRID:AB_2253745 | |
| Reelin (Reln), Mouse monoclonal G10 | Recombinant reelin amino acids 164-496, clone G10 | Sigma-Aldrich; #MAB5364 RRID:AB_2179313 | 1:1000; ICC 1:750; WB |
| S100 calcium-binding protein β (S100β), Rabbit polyclonal | S-100 isolated from cow brain | Dako A/S ( Glostrup, Denmark); #N1573 RRID:AB_2811056 | 1:500; ICC |
| Serotonin (5HT), Goat polyclonal | Serotonin | Immunostar; #20079 RRID: AB_572262 | 5000; IF |
| Stathmin 1 (Stmn1), Rabbit monoclonal | Synthetic peptide within Human Stathmin 1 aa 100 to the C-terminus | Abcam (Cambridge, MA); #ab52630 RRID:AB_2197257 | 1:7000; TSA + Citric Acid |
| SRY-Box transcription factor 10 (SOX10), Goat polyclonal | *E. coli*-derived recombinant human SOX10 | R&D Systems; #AF2864 RRID:AB_442208 | 1:75; ICC |
| Ubiquitin conjugating enzyme E2 C (Ube2c), Rabbit monoclonal | Recombinant fragment | Abcam (Cambridge, MA); #ab252940 RRID:AB_2910263 | 1:1000; TSA + Citric Acid |

*Sections underwent a permeabilization step with 100% Methanol for 20 mins at -20 °C.
**Abbreviations:** ICC, Immunocytochemistry; IF, Immunofluorescence; TSA, Tyramide Signal Amplification; WB, Western blot
