## Supplementary figures and images for "Olfactory ensheathing cells from adult female rats are hybrid glia that promote neural repair" (from another version of this preprint: version 2, 2024-09-25)

### Supplemental Figure 1

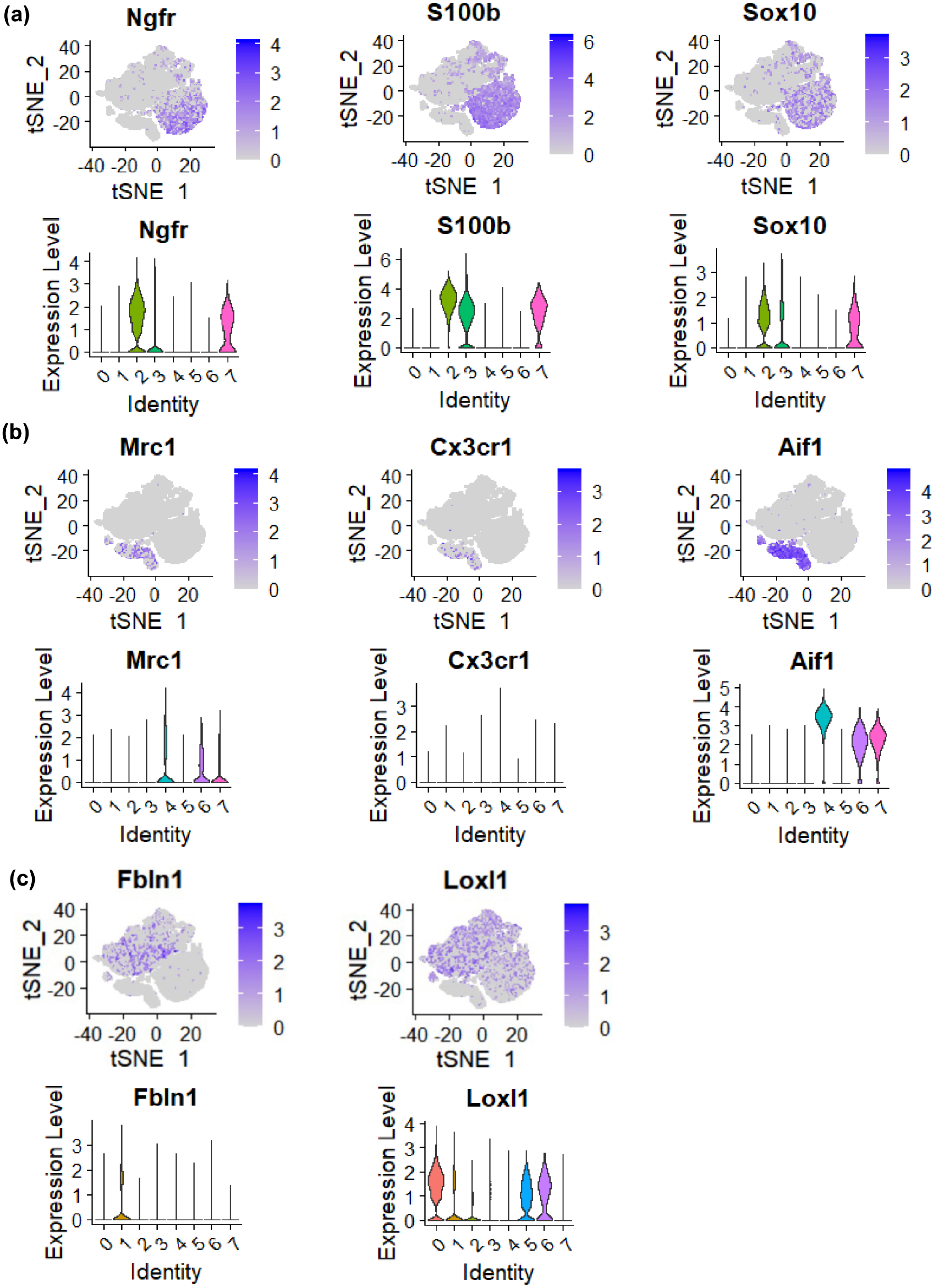

### Supplemental Figure 2

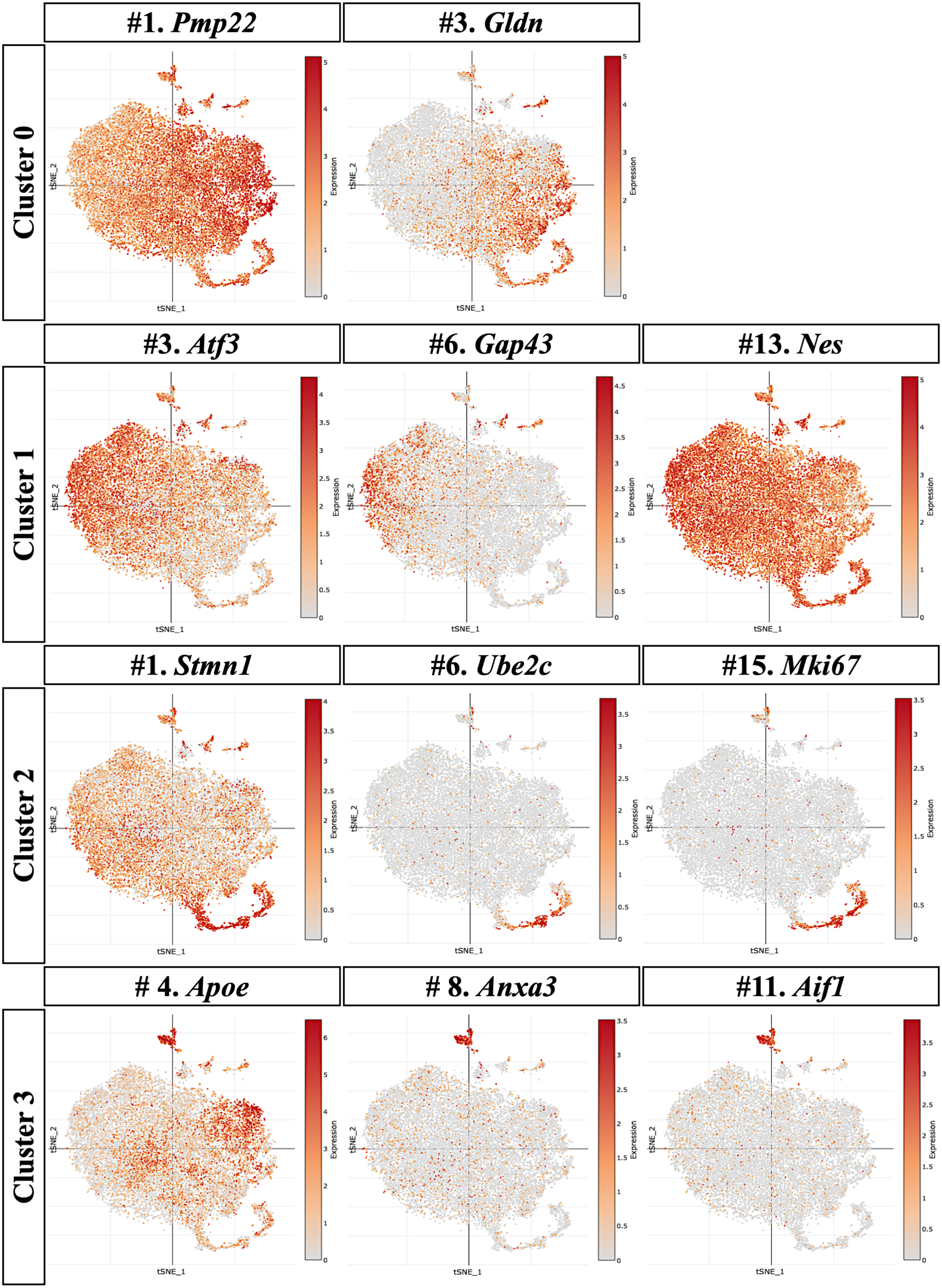

### Supplemental Figure 3

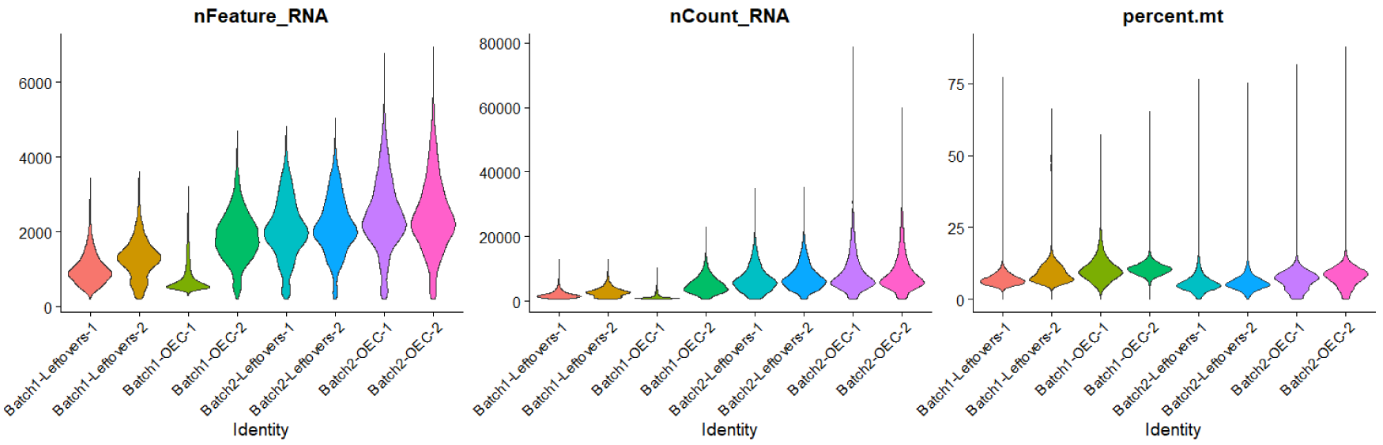

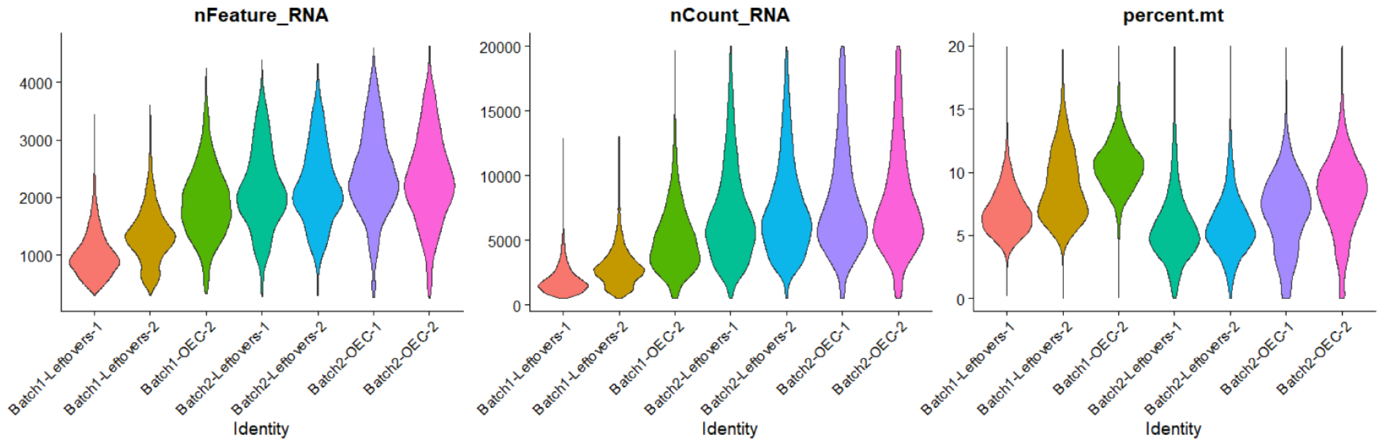
